# GRASP55 Regulates Sorting and Maturation of the Lysosomal Enzyme β-Hexosaminidase A

**DOI:** 10.1101/2024.10.16.618769

**Authors:** Sarah Reem Akaaboune, Aadil Javed, Sarah Bui, Alissa Wierenga, Yanzhuang Wang

**Author notes:** Corresponding author: Dr. Yanzhuang Wang, Institute of Neurological and Psychiatric Disorders, Shenzhen Bay Laboratory, Shenzhen, Guangdong 518132, China.

## Abstract

The Golgi apparatus plays a crucial role in the delivery of lysosomal enzymes. Golgi Reassembly Stacking Proteins, GRASP55 and GRASP65, are vital for maintaining Golgi structure and function. GRASP55 depletion results in the missorting and secretion of the lysosomal enzyme Cathepsin D (Xiang *et al*., 2013), though the mechanisms remain unclear. In this study, we conducted secretomic analyses of GRASP55 knockout (KO) cells and found a significant increase in lysosome-associated proteins in the extracellular medium. Using the lysosomal beta-hexosaminidase subunit alpha (HEXA) as a model, we found that GRASP55 depletion disrupted normal trafficking and processing of HEXA, resulting in increased secretion of the immature (pro-form) HEXA into the extracellular milieu, along with decreased levels of the mature form and enzymatic activity within the cell. GRASP55 depletion significantly reduced the complex formation between HEXA and mannose-6-phosphate (M6P) receptors (MPR), despite no overall change in MPR expression. And finally, we found there was a notable reduction in the expression of GNPTAB, leading to a reduction in M6P modification of HEXA, hindering its efficient targeting to lysosomes. These findings reveal the role of GRASP55 in regulating lysosomal enzyme dynamics, emphasizing its role in the sorting and trafficking of lysosomal proteins.

## INTRODUCTION

The organization and tethering of cisternae into stacks within the Golgi apparatus are essential for precise and sequential post-translational protein modifications, lipid synthesis, and metabolism (Zhang and Wang, 2016). Golgi Reassembly Stacking Proteins, GRASP55 and GRASP65, which display distinct localization patterns within the Golgi stack - with GRASP65 predominantly at the cis-Golgi and GRASP55 at the medial- and trans-cisternae (Barr *et al*., 1997; Shorter *et al*., 1999) - play a crucial role in the formation and organization of Golgi cisternal stacks (Xiang and Wang, 2010; Xiang *et al*., 2013). These proteins link adjacent cisternae by forming trans-oligomers, effectively tethering one cisterna to another in an orderly manner (Wang *et al*., 2003; Xiang and Wang, 2010; Jarvela and Linstedt, 2014). Depletion of GRASP proteins results in Golgi fragmentation, causing various deficiencies in post-translational protein modification, lipid metabolism, cargo sorting, and intracellular trafficking within the Golgi apparatus (Zhang and Wang, 2015, 2020).

In the Golgi, lysosomal enzymes are segregated from other glycoproteins. Most enzymes targeted to lysosomes require the acquisition of phosphomannosyl residues, specifically mannose 6-phosphate (M6P), facilitated by Golgi-resident enzymes, namely acetylglucosamine-1-phosphate transferase (GNPTAB) (Bonifacino and Traub, 2003). The binding of M6P-tagged enzymes to mannose 6-phosphate receptors is crucial for the proper sorting and targeting of lysosomal enzymes (Bajaj *et al*., 2019). Previous work has shown that in GRASP55-depleted cells, the secretion of the lysosomal enzyme cathepsin D into the extracellular space was significantly increased (Xiang *et al*., 2013). However, the precise mechanisms by which GRASP55 targets lysosomal enzymes and thus influences lysosome function are not fully understood. We, therefore, performed a secretome analysis of GRASP55 knockout (55KO) cells (Bekier *et al*., 2017; Ahat *et al*., 2022) to determine its impact on the secretion of lysosomal enzymes.

In the secretome of 55KO cells, there is an abundant secretion of lysosomal enzymes, including those involved in lysosomal storage diseases (LSDs). One such enzyme is β-Hexosaminidase A (HEXA), whose mutation results in Tay-Sachs disease. Tay-Sachs disease is a lysosomal storage disorder characterized by the progressive destruction of nerve cells in the brain and spinal cord. This disease is primarily attributed to mutations in the HEXA gene. HEX is a heterodimeric protein complex composed of α and β subunits, encoded by the HEXA gene on chromosome 15 and the HEXB gene on chromosome 5, respectively (Dersh *et al*., 2016). The α and β subunits share substantial sequence similarity, with the active site on the α-subunit efficiently hydrolyzing β-linked N-acetylglucosamine 6-sulfate-containing glycosaminoglycans and, most importantly, GM2 ganglioside. In Tay-Sachs disease, mutations in HEXA, which is normally glycosylated, prevent its transport from the endoplasmic reticulum (ER) to the Golgi (Barritt *et al*., 2017; Ramani and Parayil Sankaran, 2023). Given that HEXA was abundantly found in the secretome of GRASP55-depleted cells, we sought to investigate how GRASP protein depletion impacts the sorting and trafficking of HEXA.

This study investigates how GRASP55 regulates the targeting of the lysosomal enzyme HEXA. We found that GRASP55 depletion significantly impairs the proper targeting of HEXA to lysosomes. Specifically, 55KO cells showed a marked reduction in intracellular HEXA protein levels and activity compared to wild-type (WT) cells. Additionally, there was a notable increase in the secretion of the HEXA precursor into the extracellular medium, along with a significant reduction in the expression and maturation of GNPTAB. This was accompanied by a decrease in HEXA binding to mannose-6-phosphate receptors (MPR), which impeded its efficient lysosomal targeting. These findings reveal a mechanism by which GRASP proteins regulate lysosomal enzyme sorting and highlight the critical role of GRASP55 in ensuring proper HEXA localization and function. This may offer valuable insights into the cellular dysfunctions underlying Tay-Sachs disease.

## RESULTS

### Depletion of GRASP55 leads to an increase in the secretion of lysosomal enzymes

We conducted a comprehensive analysis of lysosomal secretion in conditioned media and lysates from WT and 55KO cells using tandem mass tag (TMT) labeling and LC– MS techniques, as previously described (Ahat *et al*., 2022). The analysis of a volcano plot comparing lysosomal enzymes secreted into the medium and those retained in WT and 55KO cells (**Figure 1A and B**) revealed significant changes (|Log2FC [55KO/WT]| > 0.5, p < 0.05). In 55KO cells, lysosomal enzymes associated with lysosomal storage diseases were notably altered. Many of these enzymes were reduced in the cell lysates, while their secretion into the medium was increased.

**Figure 1:**
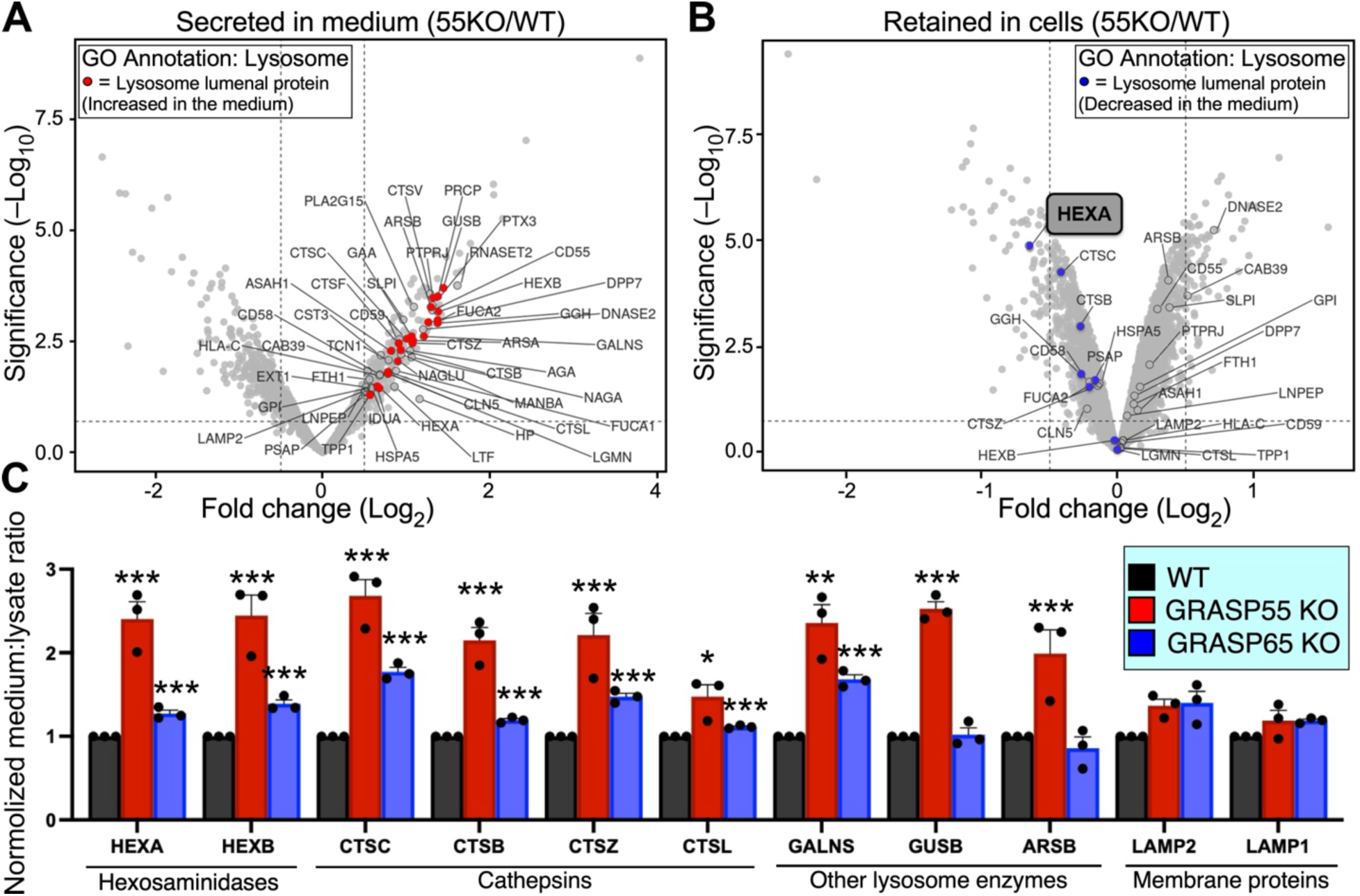
Secretome analysis of secreted proteins in 55KO cells. A) Volcano plot of secreted proteins in medium of 55KO cells versus WT secretome. Lysosomal proteins are heightened in red dots. B) Volcano plot of candidate proteins retained in 55KO cell lysates versus WT secretome. Lysosomal proteins are heightened in blue dots. HEXA is highlighted. C) Normalized medium lysates of selected lysosomal-associated enzymes.

Normalization of the ratio of lysosomal enzymes between the media and lysates of 55KO and GRASP65 knockout (65KO) cells relative to WT revealed a significant increase in lysosomal enzymes in 55KO cells, particularly HEXA and cathepsins (**Figure 1C**). These results highlight GRASP55’s pivotal role in the proper targeting of lysosomal enzymes. To gain further insights into how GRASP55 regulates the secretion and targeting of lysosomal enzymes, we focused on HEXA, a lysosomal enzyme responsible for hydrolyzing GM2 gangliosides and other molecules containing terminal N-acetyl hexosamines. Therefore, the subsequent experiments were centered on HEXA and its regulation by GRASP55.

### HEXA protein level and activity are reduced in 55KO cells

HEXA is a lysosomal enzyme synthesized in the endoplasmic reticulum (ER). It undergoes post-translational modifications in the Golgi, where it is segregated from other glycoproteins and targeted to the lysosome to perform its function. To determine whether GRASP55 regulates HEXA expression and activity in lysosomes, we conducted experiments utilizing WT, 55KO, and 55KO HeLa cells rescued by exogenous GRASP55-GFP expression (55R).

In the first set of experiments, we performed immunocytochemistry on these cells. WT, 55KO, and 55R cells were cultured, fixed, permeabilized, and immunostained with specific antibodies against HEXA. The cells were imaged using a confocal microscope, and the HEXA fluorescence level was quantified. As shown in **Figure 2A**, the fluorescence intensity of HEXA was significantly reduced in 55KO compared to WT cells. Conversely, the HEXA signal was significantly increased by GRASP55-GFP expression. Quantification of images revealed that the average fluorescence intensity of HEXA in 55KO cells (126,838 ± 51,394 SD, n = 24 cells from 6 cultured dishes) was reduced by 40% compared to that in WT cells (205,329 ± 51,995 SD, n = 24 cells from 6 culture dishes; one-way ANOVA, p<0.0001) (**Figure 2B**). In contrast, in 55R cells, HEXA intensity was significantly increased (232,120± 70,317 SD, n = 25 cells from 6 cultures) compared to that in 55KO cells and was not significantly different from WT cells (**Figure 2B**).

**Figure 2:**
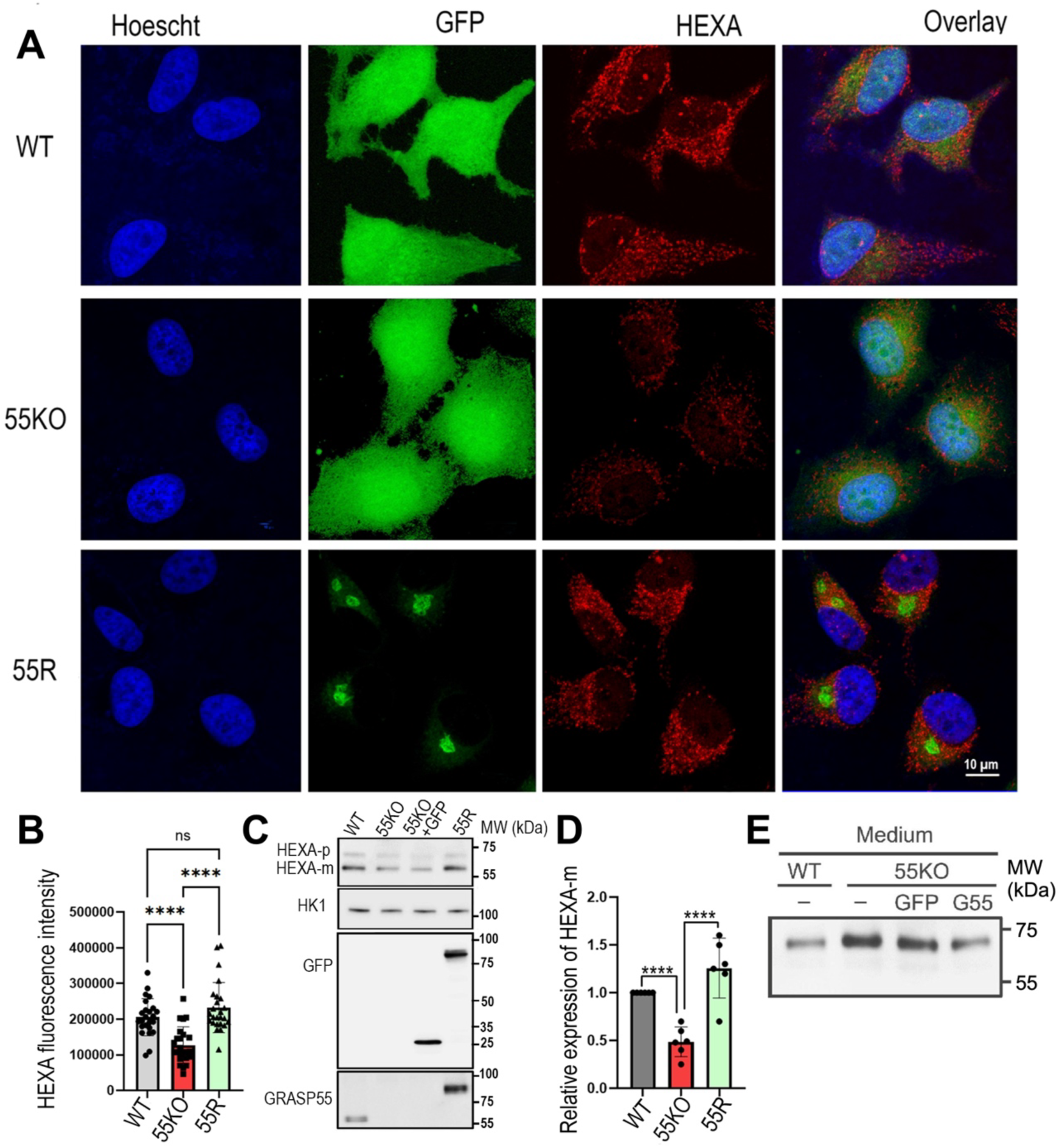
GRASP55 is essential for the targeting and maturation of HEXA in lysosomes. A) Immunostaining of endogenous HEXA protein in WT, 55KO, and 55R cells. The fluorescence signal of HEXA was significantly reduced in 55KO cells compared to WT, whereas re-expression of GRASP55 increased HEXA intensity. B) Graph summarizing the quantification of fluorescent HEXA. C) Western blot showing a notable decrease in HEXA expression levels in the lysates of 55KO cells compared to WT and 55R. D) Quantitation of mature HEXA (HEXA-m) from blot data presented in A (at least six blots were used in this quantification). D) Quantitation of the mature form of HEXA in the cell lysate. Note that the expression of HEXA-m was significantly reduced in 55KO compared to WT and 55R. E) Western blot of the secreted HEXA in the media of WT, 55KO, and 55R cells. Note that the band corresponds to the molecular weight of pro-form HEXA. The data are expressed as means ± S.D. Statistical analysis was done by one-way ANOVA (****, p < 0.0001; ns, not significant).

Western blot studies further confirmed these results. Lysates from WT, 55KO, and 55R cells were collected and subjected to immunoblotting using antibodies against HEXA. Figure 2C shows the presence of two bands corresponding to the mature (lower molecular weight) and pro-form (higher molecular weight) HEXA proteins. Quantification of the blots revealed a significant reduction (∼ 60%, ±0.3 SD, n=6) in the expression levels of the mature HEXA protein in 55KO cell lysates compared to that in WT cells. However, 55KO cells rescued with GRASP55-GFP plasmid showed a marked increase in the mature form of HEXA (**Figure 2C**). These findings suggest that GRASP55 is essential for the proper targeting, maturation, and maintenance of normal HEXA expression levels.

Because the absence of GRASP55 leads to a substantial reduction in both the pro-form and mature form of HEXA in lysates, we examined whether the pro-form of HEXA is secreted into the extracellular space instead of being targeted to the lysosomes. To test this idea directly, we conducted a secretion assay to evaluate the pro-form of HEXA secretion in the media from 55KO cells compared to WT cells and 55KO rescued cells. Medium proteins from WT, 55KO, and 55R cells were subjected to immunoblotting using antibodies against HEXA. **Figure 2E** shows a marked increase in the level of pro-form HEXA in the medium from 55KO cells. However, in 55KO cells expressing exogenous GRASP55-GFP (55R), pro-form HEXA levels were significantly reduced compared to cells lacking GRASP55. These results provide direct evidence supporting the critical role of GRASP55 in mediating the targeting of pro-form HEXA into lysosomes, thereby influencing its cellular distribution and secretion.

Finally, we examined the effect of GRASP55 on HEXA activity. We performed activity tests on the lysates and media from WT, 55KO, and 55R cells. Similar to the protein levels in cells (**Figure 3A**), the HEXA enzyme activity was markedly reduced in 55KO lysates compared to WT, with significant restoration observed in 55R cells (**Figure 3B**). Similarly, while HEXA activity in the media was considerably lower than in the lysates, it followed the same overall trends as lysate activity, with a significant reduction in 55KO compared to WT and 55R (**Figure 3C**). These results indicate that GRASP55 is essential for proper lysosome targeting and functionality for HEXA.

**Figure 3:**
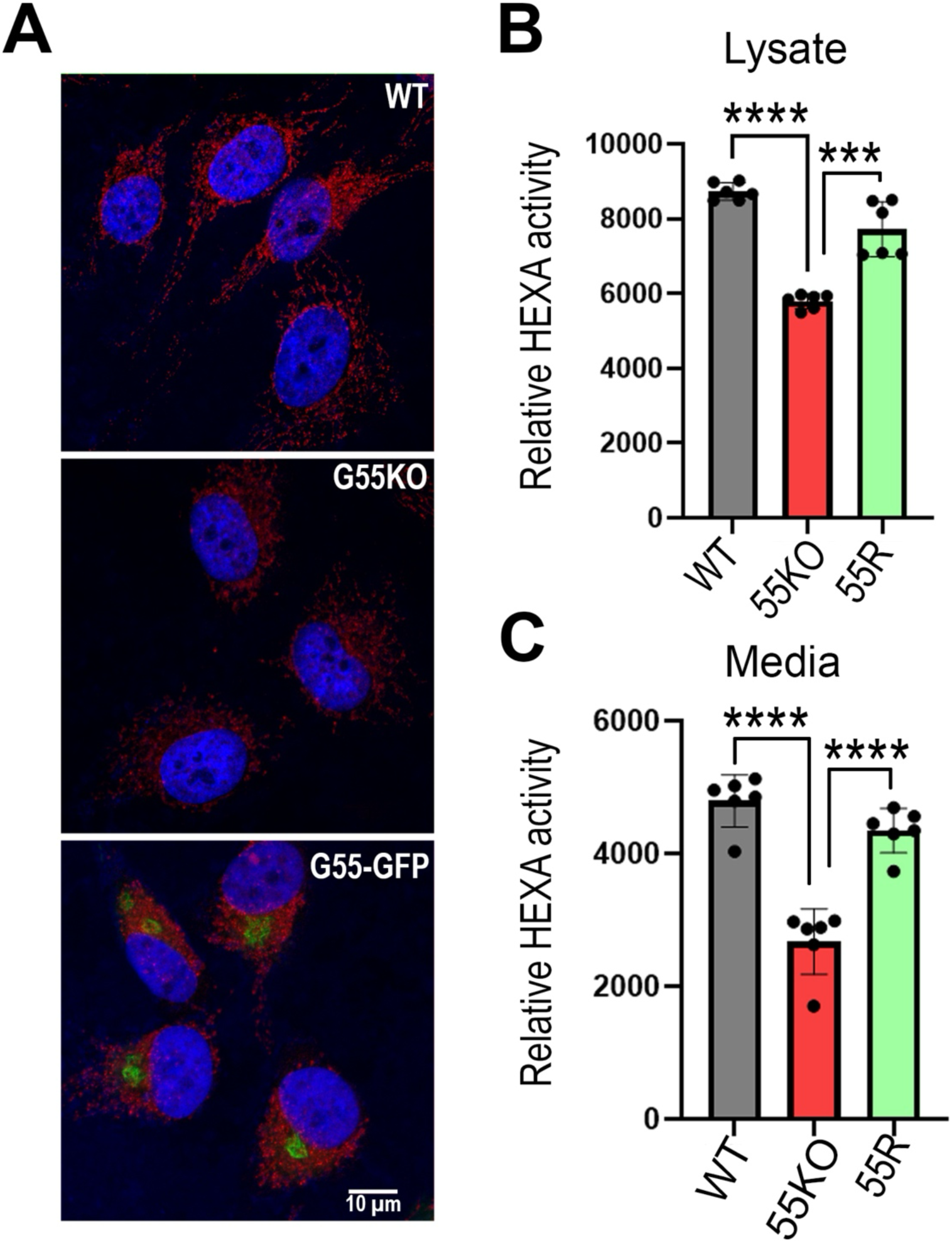
Depletion of GRASP55 induces a significant reduction in HEXA enzymatic activity. A) Images of fluorescently labeled HEXA (red) in WT, 55KO and 55R cells. B-C) Quantitation of HEXA activities in cell lysates and media. The data are expressed as means ± S.D. Statistical analysis was done by one-way ANOVA. ***, p < 0.001; ****, p < 0.0001.

### Depletion of GRASP55 has no measurable effects on the overall abundance and distribution of lysosomes

In light of the considerable increase in lysosome-associated secreted proteins in the media (Figures 1-3), we investigated whether the depletion of GRASP55 affects the number or density of lysosomes within the cells. To address this, WT, 55KO, and 55R cells were cultured for 24-48 hours, fixed, labeled with antibodies against either Lamp1 or Lamp2 (both lysosomal markers), and imaged with a Nikon ECLIPSE Ti2 Confocal microscope. **Figure 4A** shows that the overall quantity of Lamp1 appears similar across all three cell types. However, depletion of GRASP55 significantly reduced HEXA fluorescence intensity, which was restored in 55R cells. Quantification of Lamp1 fluorescence intensity within the cells revealed no significant difference between WT (186026.45 ± 92355.5 SD, n=22), 55KO (148778.9 ± 57914.58SD, n=21) and 55R cells (188346.65 ±85706.06 SD, n=32; one-way ANOVA, p=0.19) (**Figure 4B**). A similar result was obtained with Lamp2 [WT (81944.1 ±19062.4 SD, n= 11; 55KO (101060.45 ±33333.72 SD, n=13); 55R (107776.86 ± 32320.5 SD, n=26; one-way ANOVA, p=0.08) (**Figure 4B**). These results were further confirmed by western blotting, showing that the overall expression of Lamp1 or Lamp2 is similar across all types of cells (**Figure 4D**). These data suggest that the reduction of intracellular HEXA expression and activity in 55KO cells is independent of lysosomes within the cell.

**Figure 4:**
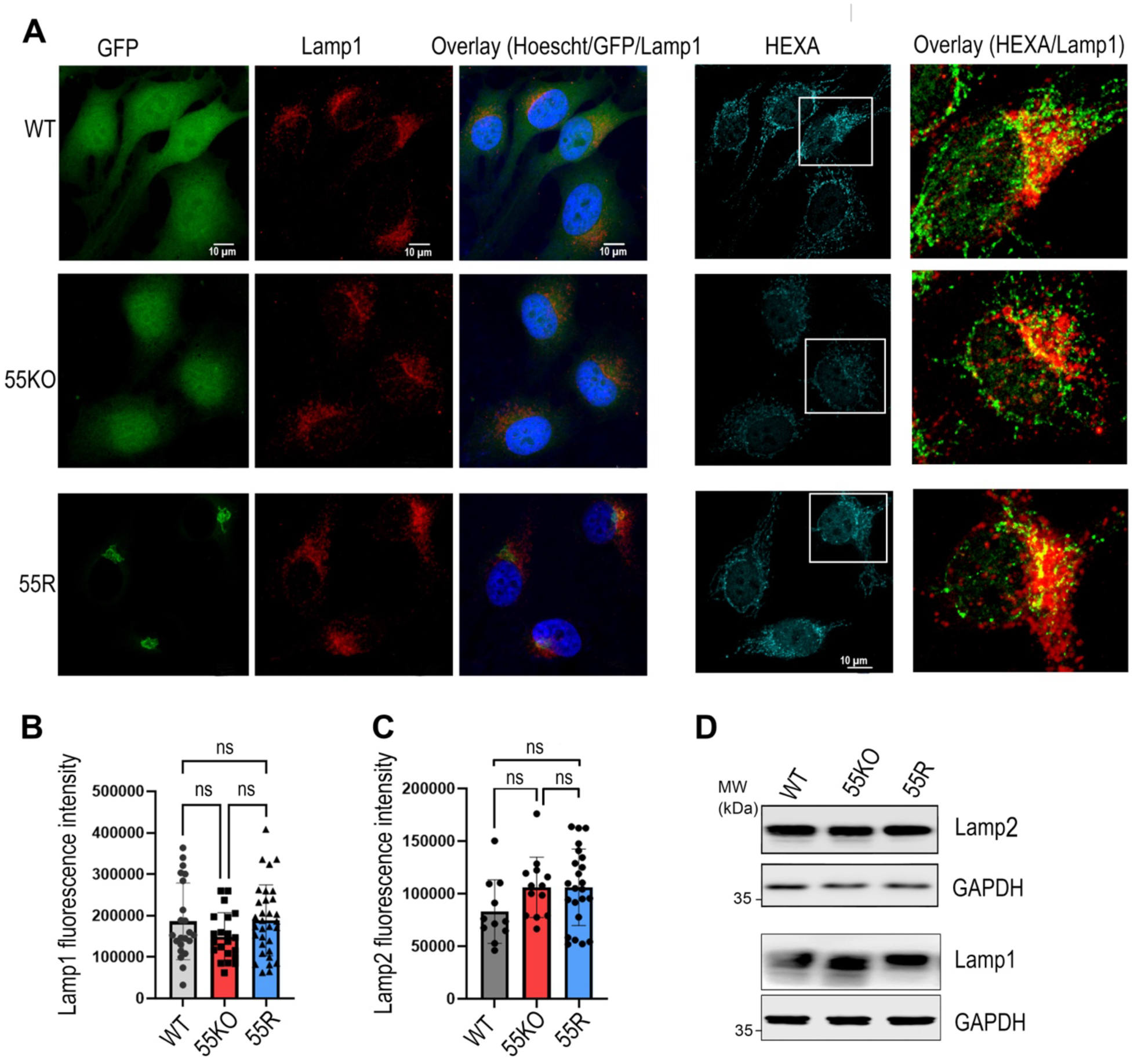
GRASP55 depletion does not affect the quantity of lysosomes despite a significant reduction in HEXA. A) Immunofluorescence images of Lamp1 in WT, 55KO, and 55R cells (left panels). Note that there are no major changes in the fluorescence intensity or localization of Lamp1 in the cell. The right panels show the overlay between Lamp1 and HEXA. Note the partial localization between HEXA and lamp1 indicated by the yellow color. B-C) quantitation of Lamp1 and Lamp2 in A. Note that there are no major changes in the fluorescence intensity of both lysosomal markers. D) Western blots of Lamp2 and Lamp1 in WT, 55KO and 55R cells. No significant changes were observed in the band intensities between the three cell lines. The data are expressed as means ± S.D. Statistical analysis was done by one-way ANOVA. Significance was defined as p < 0.05 for all analyses; ns, not significant.

### Depletion of GRASP55 impairs the binding of HEXA to the Mannose-6-P Receptor (MPR)

It is well established that in the Golgi apparatus, most lysosomal enzymes bind to mannose-6-phosphate (M6P) receptors (MPR), facilitating their targeting to lysosomes (Bajaj *et al*., 2019). Therefore, we determined whether the reduction in HEXA expression and activity is due to impaired binding to the M6P receptor or a decrease in MPR expression in 55KO cells. To distinguish between these two possibilities, we first examined whether the expression of the MPR was impaired in 55KO cells. We performed immunofluorescence microscopy on WT, 55KO, and 55R cells using specific antibodies against the cation-dependent mannose-6-phosphate receptor (CD-MPR).

Figure 5A-B shows that the fluorescent intensity of CD-MPR is not significantly different between WT (87650.31±34558.45 SD, n=12), 55KO (71504.9 ±28094.40 SD, n=14), and 55R (71920.28±19929.77SD, n=14) cells [one-way ANOVA (p=0.266)]. Similar results were obtained with the cation-independent mannose-6-phosphate receptor (CI-MPR) [WT (97430.58 ±17873.16 SD, n=12), 55KO (80022.45 ±24515.83 SD, n=11, and 55R (80990.03 ±23914.98 SD, n=12)] [one-way ANOVA (p=0.14)] (Figure 5C). These results were further confirmed by western blotting of lysates from WT, 55KO, and 55R cells using specific antibodies against both receptors. Figure 5D shows that the expression level of CD-MPR and CI-MPR in 55KO cells was not significantly different from that in WT cells. Collectively, these results indicate that HEXA reduction in 55KO cells is not due to a reduction in the amount of MPR.

**Figure 5:**
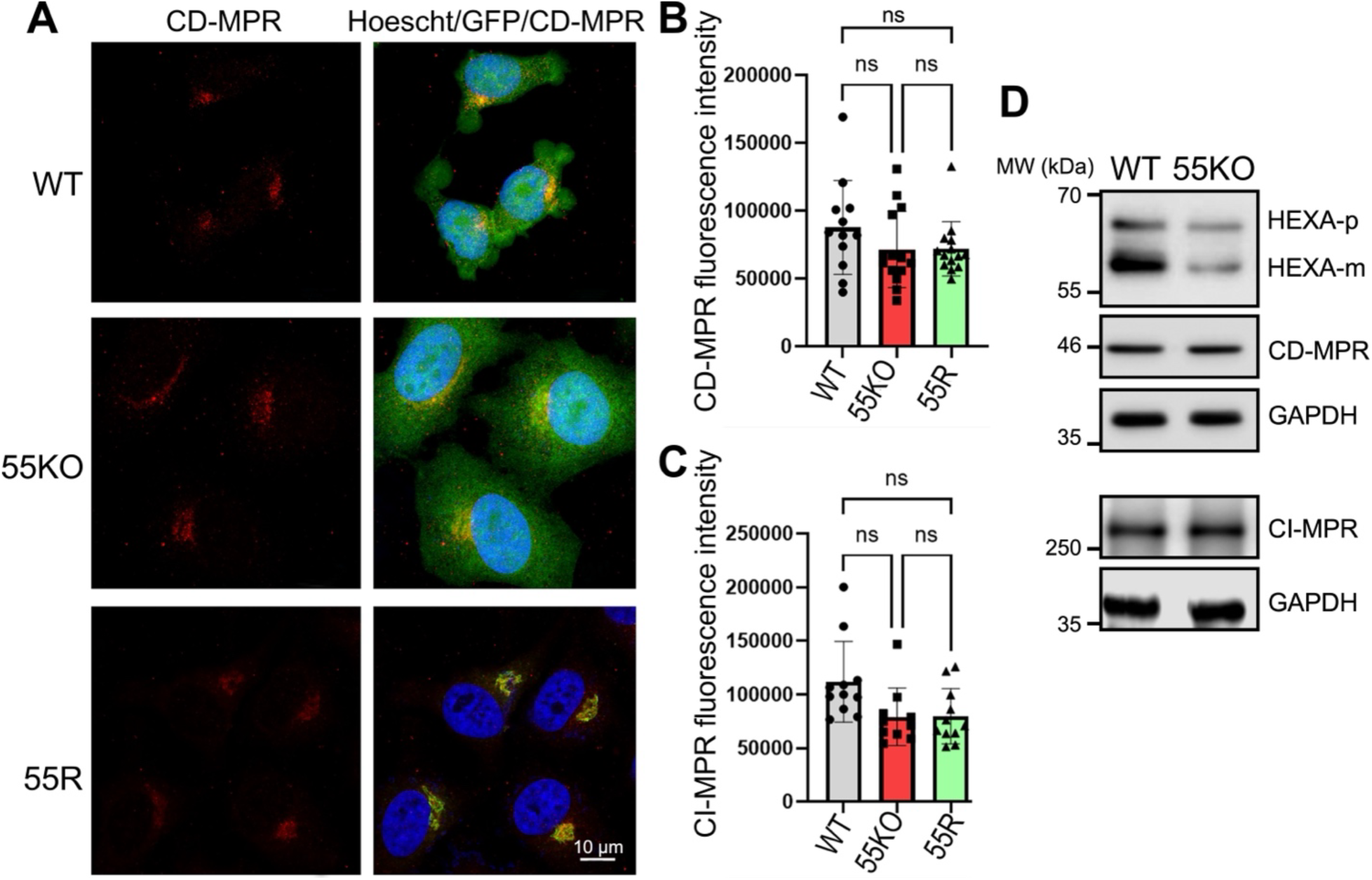
Depletion of GRASP55 has no effect on the level and distribution of mannose-6-phosphate (M6P) receptors (MPR). A) Immunofluorescence images of CD-MPR from WT, 55KO, and 55R cells. B) quantitation of CD-MPR fluorescence intensity as in A. C) quantitation of CI-MPR fluorescence intensity. Note that there are no significant changes in the cell’s fluorescence intensity or localization of CD-MPR. Data are expressed as means ± S.D. Statistical analysis was done by one-way ANOVA. Significance was defined as p < 0.05 for all analyses; ns, not significant. D) Lysates from WT and 55KO cells were subjected to immunoblotting using antibodies against CD-MPR and CI-MPR, as indicated. Note that no significant changes were observed in the MPR intensities between WT and 55KO cells.

Next, we examined the possibility that GRASP55 depletion may impair HEXA interaction with MPR. WT and 55KO cells were transfected with a flag-HEXA construct for 24 h, cell lysates were collected and subjected to immunoprecipitation using an anti-flag tag antibody, followed by western blot analysis of HEXA and CD-MPR. As expected, a significant reduction in flag-HEXA expression was observed in the cell lysate of 55KO cells compared to WT cells, indicating that loss of GRASP55 function alters the targeting of the exogenously expressed HEXA to lysosomal organelles (Figure 6A). Interestingly, when the same amount of immunoprecipitated HEXA was analyzed, the amount of co-immunoprecipitated CD-MPR was reduced in 55KO cells compared to WT (Figure 6B). This indicates that GRASP55 plays an important role in regulating the interactions between lysosomal enzymes and MPR, which controls the sorting and delivery of lysosomal hydrolases to lysosomes.

**Figure 6:**
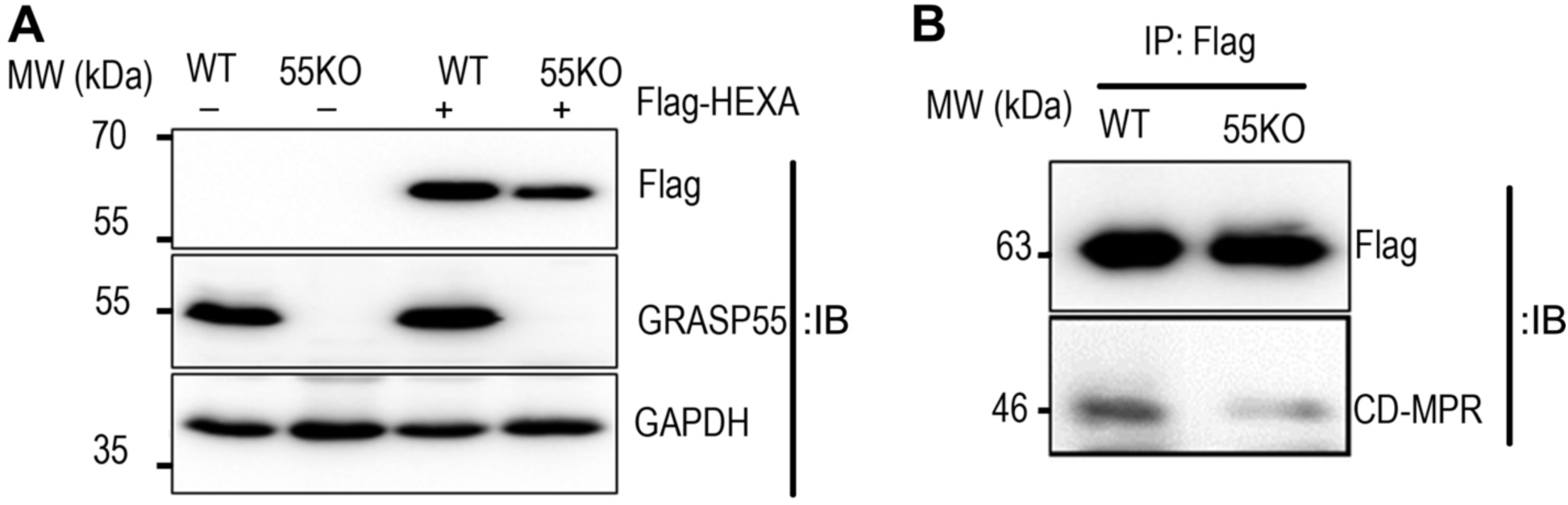
Depletion of GRASP55 reduces HEXA-MPR interaction. A) WT and 55KO cells were transfected with a Flag-HEXA plasmid, and the expression levels of mature HEXA were assessed by western blotting using antibodies against Flag. Note that the expression of Flag-HEXA was significantly reduced in 55KO cells compared to WT cells. B) Lysates from WT and 55KO cells transfected with Flag-HEXA were immunoprecipitated using an anti-Flag antibody. When equal amounts of immunoprecipitated HEXA were analyzed, co-immunoprecipitated CD-MPR levels were markedly reduced in 55KO cells compared to WT cells.

### Depletion of GRASP55 reduces the expression of N-acetylglucosamine-1-phosphate transferase (GNPTAB)

Since GRASP55 does not affect MPR expression or localization in cells (Figure 5), yet its depletion reduces HEXA levels in lysosomes, we investigated whether the absence of GRASP55 impairs HEXA sorting in the Golgi, thereby disrupting its lysosomal targeting. Given the critical role of mannose 6-phosphate (M6P) by Golgi enzymes in enabling lysosomal enzymes to bind MPRs for proper lysosomal trafficking, and the observed reduction in HEXA-MPR binding, we explored the possibility that GRASP55 regulates the lysosomal enzyme GNPTAB, which is responsible for transferring N-acetylglucosamine-1-phosphate to mannose residues on lysosomal enzymes (in this case HEXA). Due to the lack of a proper GNPTAB antibody, we used a previously established GNPTAB-HA knock-in HEK293 cell line (Zhang *et al*., 2022b) to determine the effect of GRASP55 KO on GNPTAB expression using an anti-HA antibody. Figure 7A-D shows that cells lacking GRASP55 displayed a significant reduction in the expression of the mature form of GNPTAB, along with an accumulation of the pro-form. Conversely, re-expression of GRASP55 in 55KO cells increased the level of mature GNPTAB while reducing its pro-form (Figure 7E). These findings suggest that GRASP55 regulates the expression and maturation of GNPTAB, and its reduction in the absence of GRASP55 may lead to decreased HEXA modification and reduced recognition by MPRs, thereby impairing its delivery to and activation in lysosomes.

**Figure 7:**
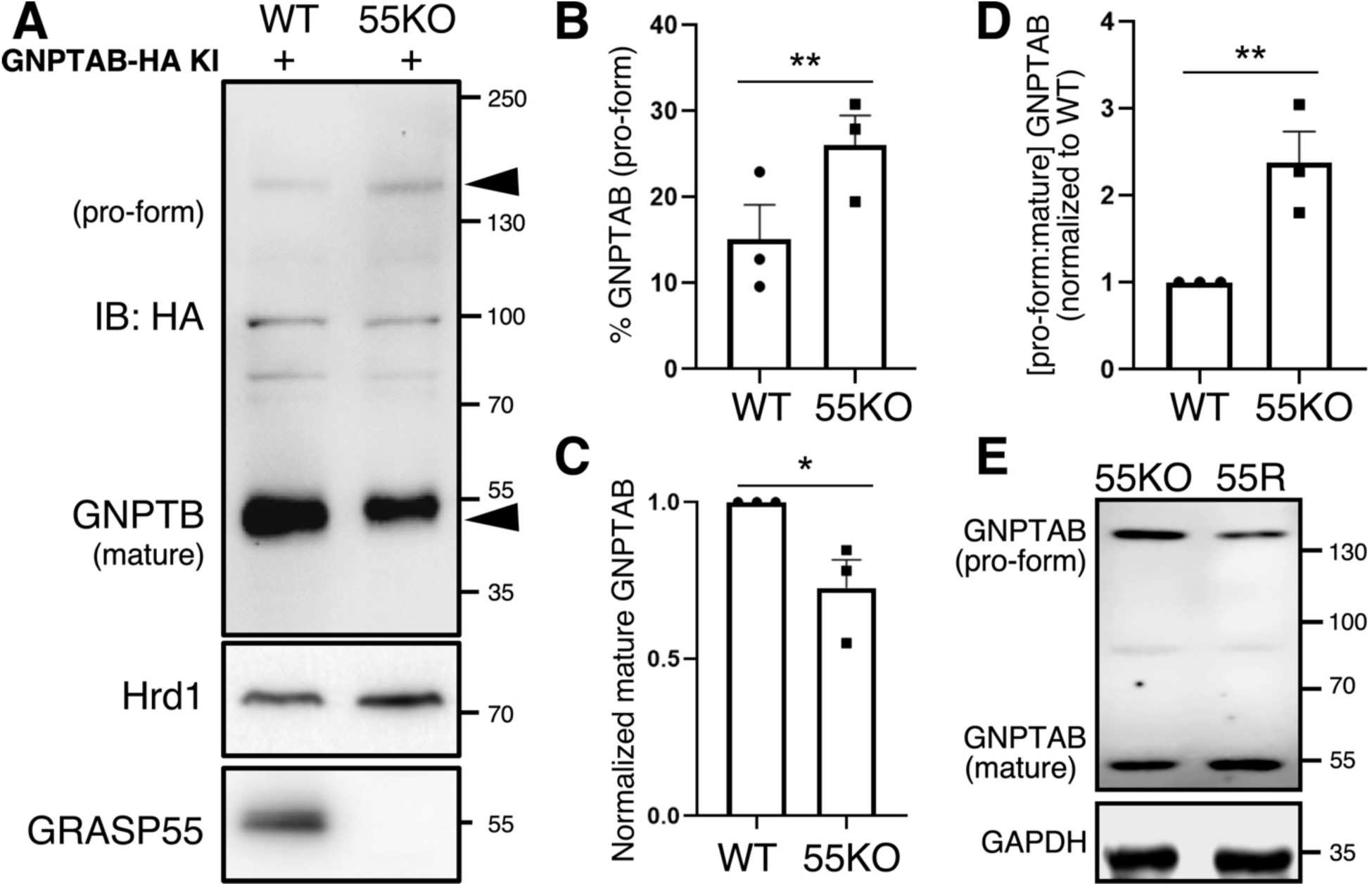
Depletion of GRASP55 results in decreased expression of the mature GNPTAB enzyme. A) Cell lysates from WT and 55KO GNPTAB-HA Knock-In HEK293T cells were probed with an HA antibody. In the absence of GRASP55, the mature form of GNPTAB-HA was reduced, while the pro-form was increased. Hrd1 was used as a loading control. B-D) Scatter plot summarizing quantification of pro-form and mature GNPTAB-HA from three independent blots (data presented as means ± S.D.). Statistical analysis was performed using Student’s t-test. *, p < 0.05; **, p < 0.01. E) Western blots of lysates from 55KO and 55R GNPTAB-HA Knock-In HEK cells using an HA antibody. In rescue cells, the mature form of GNPTAB expression increased, while the pro-form decreased compared to 55KO cells.

## DISCUSSION

This study demonstrates that GRASP55 plays a crucial role in regulating the secretion and trafficking of various lysosomal enzymatic proteins. Notably, we found that depletion of GRASP55 leads to the mistargeting of numerous lysosomal-associated enzymes into the extracellular space, along with a significant decrease in their intracellular expression. Using HEXA as a model, we show that GRASP55 depletion results in (i) a notable increase in the secretion of the enzyme’s immature form into the extracellular medium, (ii) a marked reduction in the mature form of the enzyme within the cell, and (iii) a significant decrease in its functional activity within lysosomes. Furthermore, we propose a potential mechanism by which GRASP55 controls the sorting and targeting of HEXA through the regulation of GNPTAB, a critical enzyme involved in transferring M6P onto lysosomal enzymes. In GRASP55-depleted cells, mature GNPTAB expression is substantially reduced, impairing the acquisition of M6P by HEXA. This impairment affects the enzyme’s binding to MPR, thereby hindering its efficient targeting to lysosomes. Intriguingly, despite the loss of GRASP55 function, the overall expression of MPR remains unaltered. These findings shed light on the mechanisms by which GRASP55 governs the dynamics of the lysosomal enzyme HEXA, underscoring its pivotal role in maintaining cellular homeostasis.

The secretome analysis conducted on WT and 55KO cells in this study provides valuable insights into the crucial role of GRASP55 in the targeting and secretion of lysosomal enzymes. The substantial presence of secreted lysosomal enzymes in the medium of GRASP55-depleted cells, which exhibit defects in Golgi stack formation, strongly suggests a direct link between GRASP55 function and the accurate targeting of these enzymes. This finding further supports the idea that GRASP55 plays a significant role in the subcellular localization and subsequent enzymatic activity of lysosomal enzymes within lysosomes.

It is worth mentioning that GRASP65, another Golgi stacking protein essential for Golgi stack formation and protein trafficking across the Golgi (Li *et al*., 2019), has a different impact on lysosomal enzymes. In GRASP65-depleted cells, only a few lysosomal enzymes were mistargeted and secreted into the medium (Ahat *et al*., 2022). This suggests that GRASP55 may act as a master regulator in the trafficking pathways of lysosomal enzymes and contribute significantly to lysosomal biogenesis. While both GRASP proteins are critical for Golgi stacking, their effects on the targeting and secretion of lysosomal enzymes differ. This variation may be explained by their distinct localizations and interactions within the Golgi apparatus: GRASP65 predominantly resides in the cis-Golgi, whereas GRASP55 is found in the medial- and trans-cisternae (Shorter *et al*., 1999), where sorting events occur.

The current study on the lysosomal enzyme HEXA provides insights into the mechanisms underlying GRASP55-regulated lysosomal enzyme trafficking. The absence of GRASP55 leads to a significant increase in the secretion of the pro-form of HEXA and a reduction in the mature, functional form within the cell. This raises the question of how the lack of GRASP55 causes HEXA mistargeting. Work from our lab and others has shown that GRASP55 facilitates the proper targeting of conventional and unconventional proteins to their subcellular locations via interactions with various Golgi components and trafficking machinery (Xiang *et al*., 2013). It is also worth noting that in the Golgi apparatus, the pathway of lysosomal enzymes diverges from that of other glycoproteins, with the acquisition of phosphomannosyl residues on lysosomal enzymes by the GNPTAB in the Golgi being a prerequisite for their high-affinity binding to the transmembrane mannose 6-phosphate receptors (MPR) (Staudt *et al*., 2016). The binding of phosphomannosyl-enzyme to MPRs facilitates their segregation from proteins destined for secretion and ensures their efficient targeting to lysosomes (Coutinho *et al*., 2012).

Interestingly, the current study showed that in GRASP55-depleted cells, the expression of the mature form of GNPTAB, which tags lysosomal enzymes with a M6P marker, was notably reduced. However, the overall expression of MPR remained unaltered, while the HEXA-MPR complex was significantly diminished. This strongly suggests that the reduction in mature GNPTAB levels impacts the transfer of M6P residue to HEXA, thereby reducing the amount of phosphomannosyl-HEXA. Consequently, there is a decrease in the binding of HEXA to MPR, leading to a noticeable reduction in the mature form of HEXA and its enzymatic activity in the cytosol, along with an increase in the secretion of the immature pro-form of HEXA into the extracellular medium. These findings highlight a possible mechanism in which the sequential interplay between GRASP55, GNPTAB, HEXA, and MPR in an ordered fashion regulates lysosomal enzyme sorting and trafficking. This underscores the importance of GRASP55 in maintaining proper lysosomal function.

Recent studies have shown that the transmembrane protein 251 (TMEM251), localized in the Golgi apparatus, is essential for the cleavage and function of GNPTAB, the enzyme responsible for catalyzing M6P modification. Deletion of TMEM251 results in the mistargeting of most lysosomal enzymes due to the absence of M6P modification, leading to the accumulation of undigested materials (Pechincha *et al*., 2022; Richards *et al*., 2022; Zhang *et al*., 2022b). It is possible that the significant reduction of the mature HEXA enzyme in GRASP55-depleted cells may also involve TMEM251, which warrants further investigation.

The current study provides new insights into the underlying mechanism by which GRASP55 is involved in the targeting and function of HEXA. It is worth mentioning that HEXA is an enzyme primarily responsible for breaking down a specific type of fatty substance, GM2 ganglioside, within lysosomes. In Tay-Sachs disease, a genetic mutation results in a deficiency or complete absence of functional HEXA enzyme activity, and without sufficient HEXA activity, GM2 ganglioside accumulates within lysosomes, particularly in brain and spinal cord nerve cells, leading to progressive damage to these cells, and ultimately causing neurological deterioration (Dersh *et al*., 2016). Equally important, a deficiency of GlcNAc-1-phosphotransferase results in the excessive release of lysosomal enzymes from cells, which subsequently depletes the lysosomes of multiple crucial enzymes, leading to various of lysosomal storage diseases (Khan and Tomatsu, 2020).

The significance of GRASP55 in regulating the sorting, trafficking, and activation of lysosomal enzymes highlights its essential role in maintaining cellular homeostasis and overall health. Consequently, GRASP55 could emerge as a promising therapeutic target for lysosomal storage diseases and related conditions

## MATERIAL AND METHOD

### Reagents, plasmids, and antibodies

All reagents were purchased from Sigma-Aldrich, Roche, Calbiochem, and Fisher unless otherwise stated. The following antibodies were used: Mouse monoclonal anti-HEXA (Santa Cruz, Cat#376735), Rabbit polyclonal anti-GRASP55 (Proteintech Group, Cat# 10598-1-AP), Mouse monoclonal anti-beta-Actin (Proteintech Group, Cat# 66009-1-lg), Rabbit polyclonal anti-GAPDH (Proteintech Group, Cat#10494-1-AP), Rabbit polyclonal anti-Hexokinase1 (Proteintech Group, Cat#19662-1-AP), Mouse monoclonal anti-CD-MPR (DSHB, Cat#22d4), Rabbit polyclonal anti-CI-MPR (Proteintech Group, Cat#20253), Mouse monoclonal anti-GM130 (BD Biosciences, Cat# 610823), Mouse monoclonal anti-FLAG (Sigma, Cat#M1804), Mouse monoclonal anti-GFP (Proteintech Group Cat#66002-1-lg), Rabbit polyclonal anti-HRD1 (Proteintech Group Cat#13473-1-AP), Rabbit polyclonal anti-Lamp1 (Proteintech Group, CD107a), Rabit polyclonal anti-lamp2 (Proteintech Group, CD107b), Rabit polyclonal anti-GNPTAB (SAB2106865, Millipore Sigma).

The FLAG-HEXA cDNA construct was a gift from the lab of Dr. Argon Laboratory (University of Pennsylvania), pEGFP-N1-GRASP55 WT was generated in our laboratory (Xiang and Wang, 2010). GNPTAB-HA Knock-In (KI) cell line was a gift from Dr. Ming Li’s Lab (University of Michigan) (Zhang *et al*., 2022b).

### Cell culture and maintenance

For cell culture and maintenance, WT and 55KO HeLa and HEK293T cells were cultured in Dulbecco’s Modified Eagle’s Medium (Gibco) supplemented with 10% fetal bovine serum (FBS, Hyclone), 100 units/ml penicillin and 100 μg/ml streptomycin (Invitrogen) at 37°C with 5% CO2. For transfection, cells cultured to 50-60% confluency and incubated in full growth medium for 24 h. Plasmid transfection was performed using Lipofectamine 3000 (Invitrogen) following the manufacturer’s instructions. For expression of exogenous GRASP proteins, HeLa cells of ∼60% confluency were transfected with the indicated GRASP constructs (Tang *et al*., 2010; Xiang and Wang, 2010). For a 10-cm plate, 10 µg of pEGFP-N1-GRASP65 (WT) construct or 15 µg of pEGFP-N1-GRASP55 (WT) construct was mixed with 30 µl of polyethylenimine (PEI) and 1 ml of serum-free medium for 15 min at room temperature and then added to the cells in 9 ml of DMEM containing 10% super calf serum. For the control treatment, 10 µg of pEGFP-N1 construct was mixed with 30 µl of PEI and 1 ml of serum-free medium for 15 min and then added to the cells in 9 ml of DMEM containing 10% FBS.

### Immunofluorescence, confocal microscopy, and fluorescence quantification

WT, 55KO, and 55R HeLa cells were grown on glass coverslips pre-coated with poly-lysine (Gibco) coverslips, fixed in paraformaldehyde (4% in PBS) at room temperature (15 min), washed with PBS, and permeabilized (0.2% Triton X-100 for 5 min). Cells were then incubated with 1 % Bovine Serum Albumin (BSA) prepared in PBS for 5 min on a shaker at room temperature. Cells were incubated with primary antibody diluted in 1 % BSA in PBS (1:100 for Anti-GM130, anti-HEXA, anti-Lamp1, anti-lamp2, anti-CD-MPR, anti-CI-MPR) for one hour in a humidified chamber at 37°C. Cells were washed with PBS four times and incubated with appropriate fluorescently labeled secondary antibodies for 20 min at 37°C. The secondary antibodies were either AlexaFluor 594 (anti-mouse or anti-rabbit) and AlexaFluor 488 (anti-mouse or anti-rabbit) depending on the primary antibody host (Invitrogen, Carlsbad, CA). Cells were washed with PBS four times and incubated with the Hoechst solution (1 µg/ml) for 10 min at room temperature. After the final wash, the slides were mounted with 5 µl moviol reagent, and samples were placed at 4°C overnight before imaging using a Nikon TiO2 confocal microscope. The z-stacks were then collapsed, and the contrast was adjusted using Adobe Photoshop CS2.

The fluorescence of images of cells was quantified using Fiji ImageJ (National Institutes of Health, version 1.49). Images were opened in ImageJ and converted to 8-bit binary. The cell fluorescence was then traced with the Freehand Selections area tool and four small areas of the background were traced with the Rectangle tool near the corresponding cell. These images were then measured for Area, Mean gray value, and Integrated density. The Corrected Total Cell Fluorescence (CTCF) value was calculated by using the following equation: Integrated Density – (Area of selected cell X Mean fluorescence of background). This process normalized the fluorescence intensity to account for differences in cell size and background correction.

### Immunoblotting and pull-down

WT, 55KO, and 55R cells were plated on 6 cm dishes in equal concentration and 80-90% confluency they were collected, and cell lysates were isolated as previously described (Ahat *et al*., 2022). Cells were pelleted by centrifugation and were then lysed with lysis buffer (20 mM Tris-HCl pH 8.0, 150 mM NaCl, 1% Triton X-100 supplemented with protease inhibitor cocktail (Thermo Fisher)). When cells reached, cells were washed with PBS and collected using cell scraper in PBS. Cells were centrifuged at 1200 rpm for 5 min and pelleted. The cell pellet was lysed with RIPA lysis buffer (1% Nonidet P-40, 50 mM Tris-HCl (pH 7.4), 0.25% Na-deoxycholate, and 150 mM NaCl with 1 mM NaF, 1 mM EDTA, 1 mM Na3VO4, and 1X protease inhibitor cocktail) according to the size of the pellet. The suspended pellets were kept on ice for 30 min and sonicated at 25% power for 20 seconds and then centrifuged at 12000 rpm for 10 min. The supernatant was collected as protein lysate and quantified using BCA assay. Equal amounts of protein lysates (20 µg) were run on 8-12% SDS gels and separated using electrophoresis. Subsequently, separated proteins in gels were transferred to nitrocellulose membranes using wet transfer method (Zhang *et al*., 2022a). The membranes were blocked with 5% non-fat milk solution prepared in PBST solution containing 0.1 % Tween-20 for 1 hour at room temperature. The membranes were probed with anti-Lamp2 (1:1000, ab199946), anti-Lamp1 (1:1000), anti-CI-MPR (1:2000, cation independent) (ab124767), anti-HEXA (1:500, PA579358), anti-G55 (1:2000, 10598-1-AP), anti-Flag (M2-Sigma, 1:2000), and anti-β-actin (1:2000, MA5-15739) antibodies for either 1 hour at room temperature or overnight in cold room, followed by washing with PBST for half hour with 10 minute intervals. The primary and secondary antibody dilutions were prepared in 0.5 % non-fat milk solution prepared in PBST. The primary antibodies were washed with PBST for half hour, followed by secondary antibody incubation for 1 hour at room temperature on shaker. The secondary antibodies were washed as before, followed by a 3-minute incubation with ECL reagent (BioRad or Amersham). The membranes were imaged for chemiluminescence on ProteinSimple imager (Fluorchem).

The amount of protein lysate used for each immunoprecipitation reaction was 0.5 mg with one-hour incubation performed in the anti-Flag-Agarose (40 µL) (Sigma) and used according to the manufacturer’s recommendations. After the reaction, the complexes were washed thoroughly in a RIPA-modified buffer four times before denaturation of the samples in a Laemmli buffer (2X) for 5 min at 95°C and centrifuged to obtain the eluted protein samples. The solution was then run on SDS gel for protein separation and immunoblotted for target antibodies as specified previously.

### Secretion assay

HeLa cells were seeded overnight in 35 mm plates and incubated for 4 hours in Dulbecco’s Modified Eagle’s Medium (Gibco) with 100 units/ml penicillin and 100 μg/ml streptomycin (Invitrogen) without supplementation of 10% fetal bovine serum (FBS, Hyclone), prior to the start of the experiment. 2000 μl of medium was transferred from 35 mm plates to corresponding labeled 2 ml tubes. Tubes were centrifuged at 500 rcf for 10 min, after which 1800 μl of medium was removed and transferred to a new labeled corresponding 2 ml tubes. Tubes were then centrifuged at 400 rcf for 15 min, after which 1600 μl of medium was removed and transferred to new labeled corresponding 2ml tubes. Tubes were then centrifuged at 20,000 rcf for 10 min, after which 1400 μl of medium was removed and transferred to new labeled corresponding 2ml tubes. Then, 2.2 g of Trichloroacetic Acid (Thermo Fisher, A11156.30) was dissolved in 1 mL of Mili-Q water in a 15 mL conical tube to create a 100% Trichloroacetic Acid (TCA) solution. 154 μl of 100% TCA solution was transferred to each corresponding 2ml tube containing 1400 μl of medium. Tubes were incubated on ice for 10 min. Then, 10% TCA solution was made, by diluting 1 ml of 100% TCA solution in 9 ml of Mili-Q water in a 15 ml conical tube. 500 μl of 10% TCA solution was added to tubes. Tubes were incubated in ice for 20 min. Tubes were then centrifuged at 20,000 rcf for 30 min. Supernatant from tubes was carefully aspirated and 500 μl of ice-cold acetone was added. Tubes were gently rocked back and forth and centrifuged for an additional 20,000 rcf for 10 min. Acetone was aspirated from tubes and tubes were inverted for 3 min to reveal a cloudy pellet. 28 μl of 2X SDS-PAGE sample buffer was added to the tube and manually interspersed with the pellet to create an evenly distributed sample for western blotting.

### HEXA activity assay

HeLa cells were seeded overnight in 35 mm dishes. Media was aspirated and cells were washed with cold PBS. Cells were scraped, transferred to 1.5ml tubes and resuspended in 1 ml of PBS, and centrifuged at 1000 rpm for 5 min. Supernatant was aspirated from tubes to reveal a pellet. Pellet was resuspended with lysis buffer in amounts correlated to approximately double the pellet size, and then incubated on ice for 10 min. Tubes were centrifuged for 5 min at 15,000 rpm and lysate was subsequently isolated and quantified using a Bradford Kit (BioRad). Volume required for 5 µg of lysate was calculated and topped up to a total volume of 50 μl using lysis buffer. Lysates from WT and KO were incubated for 1-2 hours in the dark at 37 degrees with methylumbelliferyl-N-acetyl-β-D-glucosaminide (4-MUF-NAG), a synthetic uncharged fluorogenic substrate for HEXA. Mixtures were read on a TECAN plate reader at excitation and emission 328/460 nm. Changes in fluorescence were reported as a measure of HEXA activity. Cell lines were plated in 12-wells plate and using the β-hexosaminidase assay kit (CS0780; Sigma). Briefly, the β-hexosaminidase activity was determined by the addition of 90 μl of assay buffer [0.09 M citrate buffer (pH4.8); 1 mM 4-nitrophenyl N-acetyl-β-D-glucosaminide] to 10 μl of cell lysates. the reaction mixture was incubated at 37°C for 10 min and stopped by the addition of 200 μl of stop solution (0.5 N of NaOH). Fluorescence was measured (Ex/Em = 328/460 nm). Fold increases in protease activity were determined by comparing the relative fluorescence units (RFUs) against the levels of the controls and visualized using a Tecan D300e.

### Quantification and statistics

All western blots were quantified using FiJi ImageJ. Background values are subtracted from raw values, and all values are normalized to β-actin expression within the sample for each biological replicate unless otherwise specified in figure captions. All graphs were made in Prism GraphPad 9.0. Statistical analyses were performed using a two-tailed Student’s t-test or one-way ANOVA. Differences in means were considered as no statistical significance (n.s.) if p ≥0.05. Significance levels are as follows: *, p < 0.05; **, p < 0.01; ***, p < 0.001, ****, p < 0.0001.

## ACKNOWLEDGMENTS

We thank all members of the Wang Lab, especially Dr. Erpan Ahat, Dr. Jianchao Zhang, and Jason Kwon for assistance with experiments and stimulating discussions. We also thank Dr. Yair Aron (University of Pennsylvania) for the FLAG-HEXA construct and Dr. Ming Li (University of Michigan) for the GNPTAB-HA knock-in HEK cell lines. This work was supported by National Institutes of Health (Grant R35GM130331) to YW. SA was an undergraduate at the University of Michigan whose research is supported by National Institutes of Health (Grant 3R35GM130331-05S1).

## AUTHOR CONTRIBUTIONS

S.R.A, S.B., and W.Y. conceptualized the study. S.R.A, A.J., and S.B. performed experiments. S.R.A wrote the first version of the manuscript and interpreted the data. W.Y. edited the manuscript and interpreted the data in the paper. A.W. analyzed the data. All authors read and approved the final version of the manuscript.

